# Optimization of extraction, Structural characterization of immunomodulatory acidic polysaccharide from *Euphorbia caducifolia*

**DOI:** 10.1101/2023.03.10.532010

**Authors:** Kusuma Venumadhav, Kottapalli Seshagirirao

## Abstract

Heating confluence method in combination with response surface methodology (RSM) was performed to optimize polysaccharides extraction from *Euphorbia caducifolia* Haines. The Box-Behnken Design was used to optimize the extraction conditions with three parameters such as temperature (A), Solid to liquid ratio (B), and time (C). The percentage yields obtained were fitted to a 2^nd^ order polynomial equation using multiple regression analysis. From the regression equation, the optimal polysaccharides extraction parameters for 2.3% of yield are Temperature-120°C, Solid to liquid ratio-75 g/L, and Time-2 h. Further, crude polysaccharide was purified and disclosed GlaUA, Rha, Glc, Gal, Xyl and Ara (1:2.5:4:3.6:3.7:3.3) ratio of poysaccharide. Thermogram of the EC polysaccharide showed typical natural polysaccharide behavior, and hydrophilic nature of polysaccharide attributed to endothermic water loss. Furthermore, purified polysaccharides exhibited strong *in vitro* free radical scavenging activities, reducing ability, and immunomodulatory activity by the effective production of pro-inflammatory cytokines with RAW 264.7 cells.

## 1. Introduction

Polysaccharides can serve as an energy source and structural components to the cell with its highly complex structure (Wang, Luo, & Ena, 2007). Polysaccharides from higher plants show significant therapeutic properties by its non-toxic nature and ability to not cause any side effects compared to bacterial and fungal polysaccharides. Thus, botanical polysaccharides are ideal therapeutics for immunomodulatory, antitumor and wound healing activities (Schepetkin & Quinn, 2006).

*Euphorbia caducifolia* Haines, a Euphorbiaceae species grows extensively on rocky areas of arid and semi-arid regions with extensive branching and clumps of phylloclades (Shaik, Valli, Rajeswari, & Kumar, 2013). Phylloclades are fleshy, succulent with spiny stipules. Milky and sticky natured latex flows through laticiferous cells along with proteins, phytosterols and terpenoids (Afza, Khan, Malik, & Badar, 1989). Ethnomedical uses of the plant were reported, latex for wound healing activity and other skin infections (Goyal, Nagori, & Sasmal, 2012), root extracts for snakebite (Chavre B.W, 2013), leaves for antimicrobial activity (Kapoor, Mishra, Acharya, Lakhera, & Purohit, 2013) and phylloclades as biofuels (Shaik et al., 2013).

Response Surface Methodology (RSM) is known to be the best technique to evaluate the relationship between parameters and the yield (D. Ye et al., 2015). Extraction temperature, solid to liquid ratio and time are the common variables which affect the yield. In RSM, Box-Behnken experimental design is more efficient and easier method to analyze and interpret the variables affect the yield (Chaiklahan et al., 2013).

Novel immunomodulatory polysaccharides from the unexplored arid plant exhibit the wound healing activity. This study conducted to evaluate and optimize the variables affecting the novel polysaccharide yield from the arid plant using a single factorial (RSM), three level-three variable design (BBD). In addition, reporting its structural and biological studies as therapeutic interests.

## 2. Materials and Methods

### 2.1. Plant material

*Euphorbia caducifolia* was collected from hills of the bongiri village, Yadadri District, Telangana State, India. The plant specimen was authenticated and deposited at University of Hyderabad Herbarium (Voucher No.UH2027). The collected stems were dried at room temperature until it gains constant weight. Dried stems were ground to pass 2 mm sieve and proceeded for the extraction.

### 2.2. Extraction process

The powder was freed from oils, fats, fatty acids, and other phytochemicals by sequential extraction using the soxhlet apparatus with hexane, chloroform and methanol. The residual marc was dried and used for polysaccharides extraction (Huang et al., 2010).

#### 2.2.1. Water extraction

Response surface methodology was used to extract crude polysaccharides in coercion with hot water method. In brief, dried marc was suspended in established volume of distilled water at designed temperature and time. The suspension was centrifuged, and collected supernatant was concentrated under vacuum at 45°C. The polysaccharides from the supernatant was precipitated with absolute ethanol and garnered the pellet by centrifugation. The precipitated pellet was washed with different percentages of ethanol to remove soluble material and deproteinized using sevag method (Zou, Jiang, & Tian, 2015). A brown coloured crude polysaccharide was obtained (C.-L. Ye & Lai, 2015) and the concentration was determined by using Phenol-Sulphuric acid method (DuBois, Gilles, Hamilton, Rebers, & Smith, 1956). The yield was calculated using the following equation:

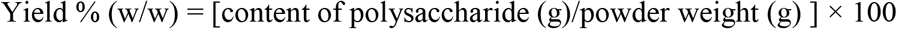

#### 2.2.2. Response Surface Methodology (RSM) Experimental Design

Box-Behnken design (BBD), a three-level three-factor design was used to optimize the polysaccharides extraction. Parameters like temperature (A), solid to liquid ratio (B) and time (C) were the independent variables, and the polysaccharide yield was taken as response function (R) to the independent variables. The BBD matrix consisted of total 17 experimental points in random order with five replications of center points (D. Ye et al., 2015). Each variable was set at three levels and coded −1, 0, +1 for low, intermediate and high values respectively (Table 1). The fitted polynomial equation was expressed as surface and contour plots for visualizing the relationship between three independent variables and the response variable(Yuan, Zeng, Nie, Luo, & Wang, 2015).

**Table 1.** Factors and levels in Box-Behnken Design used to analyze extraction yield of polysaccharides from *Euphorbia caducifolia*.

| Variables for Extraction | Factor Level |  |  |
| --- | --- | --- | --- |
|  | -1 | 0 | 1 |
| A: Temperature (°C) | 40 | 80 | 120 |
| B: Solid to liquid ratio (g/L) | 25 | 50 | 75 |
| C: Time (h) | 1 | 2 | 3 |

### 2.3. Purification and structural characterization

#### 2.3.1 Purification

Crude polysaccharide obtained was redissolved in water, and applied on DEAE Cellulose-52 column (2.5×30) and equilibrated with distilled water. The polysaccharides were eluted with stepwise NaCl solution (0, 0.1, 0.5 and 1M). The eluates containing polysaccharides was pooled, concentrated and dialyzed against distilled water. Dialyzed fraction was lyophilized (considered as EC polysaccharide or ECP) and stored at −20°C (H. Ye, Wang, Zhou, Liu, & Zeng, 2008). UV-Visible spectrum was measured for purity of the sample, polysaccharides dissolved in distilled water and spectrum recorded at a frequency range 1100-190 nm (Yuan et al., 2015).

#### 2.3.2 Structural characterization

##### 2.3.2.1 Monosaccharide analysis

EC polysaccharide was analyzed for monosaccharide composition using PMP derivitization (Yang, Zhao, Wang, Wang, & Mei, 2005). The polysaccharide was hydrolyzed in 2.5 M TFA for three h at 121°C in a sealed tube, and residual acid was removed with methanol under N2 stream. The residue was dissolved in water. Hydrolysed solution or monosaccharide mixture (100μl) was mixed with 0.3M NaOH (100μl), and methanolic PMP (0.5M, 100μl), incubated for 100 min at 70°C. Reaction mixture was cooled and neutralized with 0.3M HCl. Resultant solution was washed with chloroform in three successive orders and the final aqueous layer was analyzed by HPLC.

##### 2.3.2.2 Functional group analysis

ATR-IR was used to analyse the functional groups present in the polysaccahrides. Polysaccharide structure like glycosidic linkage, monosaccharide type and functional group attached could be investigated using IR spectroscopy. 5 mg of polysaccharide was anlysed by placing in slot of Bruker Alpha ATR-IR and measured spectrum.

##### 2.3.2.3 Linkage Analysis

Methylation studies used to determine the monosaccharide linkage in the polysaccharide (A Pettolino, Cherie, B Fincher, & Antony, 2012). Hydroxyl groups of the EC polysaccharide was methylated using methyl iodide in the presence of alkaline DMSO for 40 min, and reaction ceased with the addition of water. Extracted the methylated polysaccharide with DCM, and hydrolyzed with TFA and analyzed for linkages injecting into GC-MS.

### 2.4. Thermal Analysis (TG-DTA)

Mettler TGA/DSC1 was used to determine the thermal properties of polysaccharides.

EC polysaccharide was weighed to 4.1 mg and heated at a rate of 10°C/min from 40°C −600°C under the flow of N_2_ atmosphere (Janarthanan, Zin Wan Yunus, & Ahmad, 2003).

### 2.5. Antioxidant activity

#### 2.5.1. DPPH-free radical scavenging activity

Blois method was used to determine the free radical scavenging activity of EC polysaccharide using DPPH radical with minor modifications (Molyneux, 2004). 1 ml of DPPH (0.1mM) was added to 3ml of various concentrations of polysaccharides (0.125-2 mg/mL) and incubated for 30 min in the dark and absorbance was measured at 517 nm. Ascorbic acid was used as a standard. Three separate experiments were performed, and the percentage inhibition of the polysaccharides was calculated by the following equation: (Kumar et al., 2005)

Percentage inhibition = [(A control – A sample)/A control] × 100

Where A- Absorbance.

#### 2.5.2. Nitric Oxide radical scavenging activity

Sodium Nitroprusside gives nitric oxide (NO^·^) in aqueous solution at pH 7.2-7.4, which reacts with oxygen to produce nitrite ions and the formed nitrite ions determined by Griess reagent (Griess diazotization reaction). 3 ml reaction mixture containing 10mM Sodium Nitroprusside in phosphate buffered saline pH 7.2 and EC polysaccharide with different concentrations (0.125-2 mg/mL) were incubated at 25°C for 2 h. 0.5 ml aliquot of reaction mixture was added to 0.5 ml of Griess reagent (equal parts of 1% sulphanilamide in 5% H_3_PO_4_ and 0.1% naphthyl ethylenediamine dihydrochloride), and the absorbance was measured at 546nm. BHT (Butylated hydroxytoluene)used as reference compound. All three experiments were done separately, and the percentage inhibition was calculated by comparing control and test samples (Green et al., 1982).

#### 2.5.3. Reducing power

1 ml of EC polysaccharide (0.125-2 mg/mL) was added to 2.5 mL of phosphate buffer (pH 6.6, 0.2 M) and 2.5 ml of 1% potassium ferricyanide [K_3_Fe(CN)_6_] and incubated at 50°C for 30 min. 2.5 ml of 10% TCA was added to the mixture and centrifuged for 10 min at 3000 rpm. To the 2.5 ml of supernatant, 2.5 ml of distilled water and 0.5 ml of 0.1% FeCl_3_ added. BHT used as reference compound. The absorbance was measured at 700 nm. The increase in the absorbance indicates the high reducing power (Yildirim, Oktay, & Bilaloglu, 2001).

### 2.6. Emulsifying activity

The emulsifying activity of EC polysaccharide was performed by turbidimetry method (Benhura & Chidewe, 2004). 3ml of different concentrations of EC polysaccharide in water was added to the 1 ml of vegetable oil (castor oil) and homogenized for 10 min. A 50 μl aliquot was diluted with 5ml of 0.1% Sodium dodecyl sulfate, and the absorbance was measured at 500 nm. The ability of the emulsion was determined by recording the absorbance at 0 h, and the stability of the emulsion was determined by allowing the emulsion at room temperature for 24 h. The turbidity of the emulsion was calculated by:

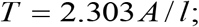

where T-Turbidity; A-Absorbance; *l*-Path length of the cuvette;

### 2.7. MTT Assay

Murine macrophages (RAW 264.7) was used for the experiment. The cells (1×10^4^) were inoculated in 96 well plate and incubated for 24 h in an incubator (5% CO_2_, 37°C). The cells were treated with different concentrations EC polysaccharide and incubated for 24 h. After incubation, 3(4,5-dimethyl thiazol-2-yl)-2,5-diphenyl tetrazolium bromide (MTT) was added as 5 mg/mL and incubated further for 4 h. 100 μl of DMSO added to the wells and absorbance taken at 590nm. Percentage of cell viability was calculated with untreated cells as a control (Khan et al., 2006).

### 2.8. Cytokine Measurement

RAW 264.7 cells (5 × 10^5^ cells/mL) were stimulated with EC polysaccharide and the levels of cytokines, Tumor necrotic factor –α (TNF-α) and Interleukin-6 (IL-6) were measured after 24 h of incubation (Schepetkin et al., 2008; Xu et al., 2006). TNF-α and IL-6 concentrations were determined by extrapolation from the standard curve, according to the manufacturer’s protocol (BD Bioscience).

## 3. Results and Discussions

### 3.1. Polysaccharides extraction

Response surface methodology, a statistical technique used to analyze the relationship between parameters and the yield (D. Ye et al., 2015). The polysaccharide yield obtained with an experimental matrix of the Box-Behnken design were calculated and showed the yield in the range of 0.9% to 2.4% (Table 2). As single factor analysis, temperature played a significant role in the percentage yield of the polysaccharides compared with solid to liquid ratio and time by enhancing diffusion coefficient of the polysaccharide at high temperatures (Ros et al., 2004). High temperatures (>90°C) were used to extract polysaccharides from algae and plant sources (Chaiklahan et al., 2013). The response surface curves (3D) and contour plots (2D) of the design gives a visual relation between parameters to the percentage of yield (Fig.1). Although the rise in temperature increases the polysaccharides yield, its interaction with solid to liquid ratio and time also affect the yield percentage. The influence of temperature on the yield with solid to liquid ratio shows statistical significance but not with time, albeit longer time affect the yield (Supplementary Table 1). At 120°C, 2.36% of yield was obtained with 50 g/L of solid to liquid ratio and 3 h of time whereas 2.30% of yield was obtained with 75 g/L of solid to liquid ratio and 2 h of time, and the insignificant difference of the yield (0.06%) shows the interchangeable effect of the other two parameters on the yield. From RSM results it observed that yield saturation reached to 1.5% & 2.4 % at below and above 90°C respectively. The interactions between the variables and yields were related with the help of multiple regression analysis by following second-order polynomial equation:

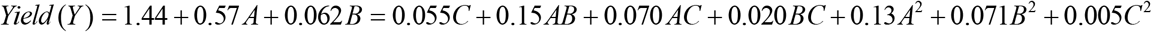

**Fig. 1.**
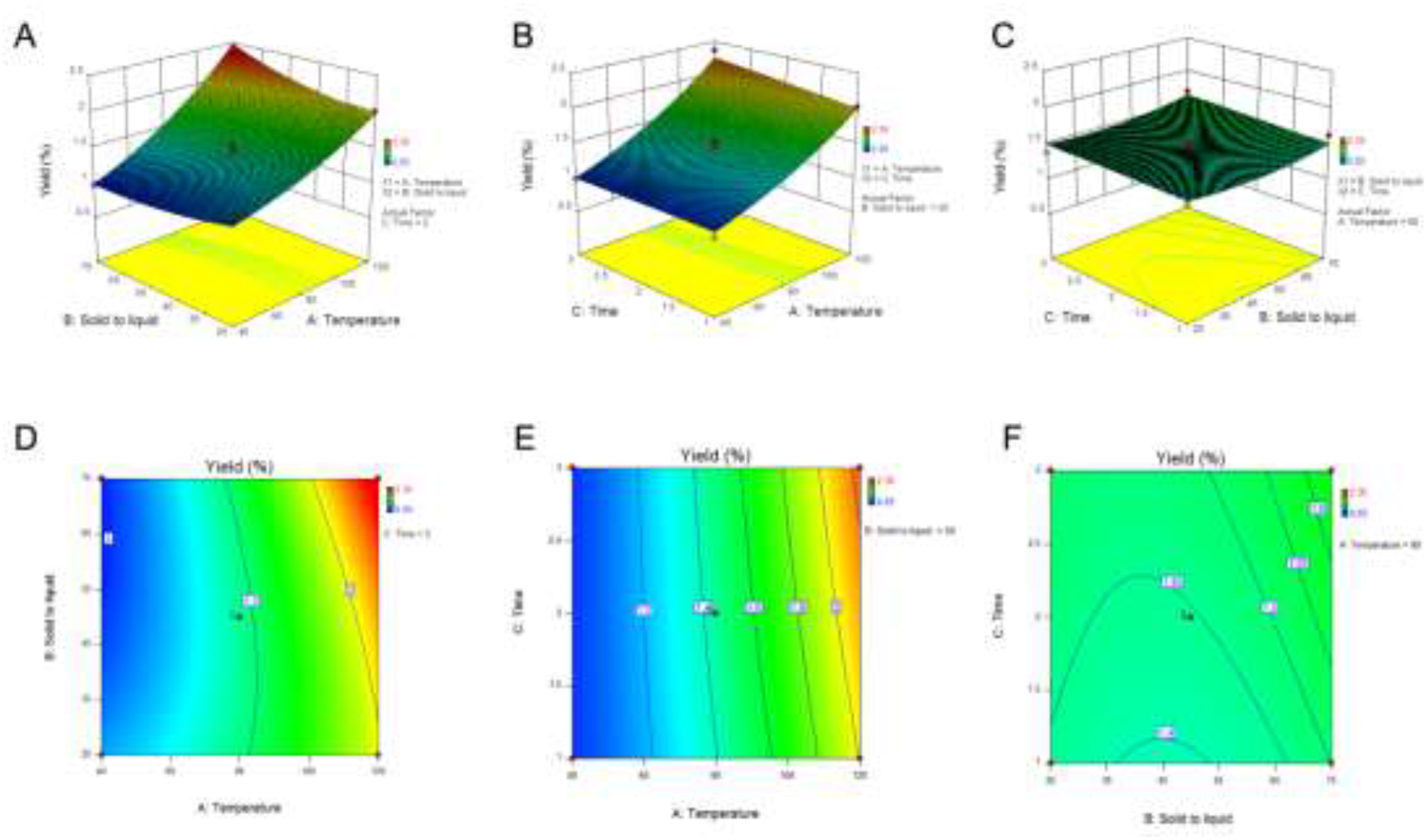
Response surface curve (3D) showing the relation between parameters and polysaccharide yield; A) temperature and solid to liquid ratio, B) temperature and time, C) solid to liquid ratio and time. Contour plots (2D) showing the relation between parameters and polysaccharide yield; D) temperature and solid to liquid ratio, E) temperature and time, F) solid to liquid ratio and time.

**Table 2.** The Box-Behnken Design matrix and results of extraction yield (%) of polysaccharides from *Euphorbia caducifolia*

| Run | A:<br>Temperature<br>(°C) | B:<br>Solid to liquid ratio<br>(g/L) | C:<br>Time<br>(h) | Yield<br>(%) |
| --- | --- | --- | --- | --- |
| 1 | 80 | 25 | 1 | 1.39 |
| 2 | 80 | 75 | 1 | 1.62 |
| 3 | 80 | 75 | 3 | 1.69 |
| 4 | 40 | 50 | 1 | 0.93 |
| 5 | 80 | 25 | 3 | 1.38 |
| 6 | 40 | 75 | 2 | 0.96 |
| 7 | 80 | 50 | 2 | 1.46 |
| 8 | 80 | 50 | 2 | 1.53 |
| 9 | 80 | 50 | 2 | 1.40 |
| 10 | 40 | 25 | 2 | 1.29 |
| 11 | 120 | 75 | 2 | 2.30 |
| 12 | 40 | 50 | 3 | 0.98 |
| 13 | 80 | 50 | 2 | 1.40 |
| 14 | 80 | 50 | 2 | 1.43 |
| 15 | 120 | 50 | 3 | 2.36 |
| 16 | 120 | 25 | 2 | 2.01 |
| 17 | 120 | 50 | 1 | 2.03 |

From the regression analysis, the high value of the determination coefficient R^2^ (0.97) and the adjusted R (0.97) explains the high fitness of the regression model and correlation between the values. The Analysis of Variance (ANOVA) of the model (Supplementary Table 1) suggests that the temperature variable affects the yield significantly as well as its interaction with solid to liquid ratio variable influence the yield (P<0.05). The optimum conditions for the better extraction of the polysaccharides are 120°C of temperature, 75 g/L of solid to liquid ratio and 2 hr of time which correlates with predicted yield (2.425%) and experimental yield (2.30%).

### 3.2. Purification

The lyophilized crude polysaccharide was chromatographed on an anion exchange column to get acidic polysaccharide fraction. Use of an anion exchange chromatography to elute the acidic polysaccharides has been reported from wheat, sugar beet by Neukom et al. (1960) (Jermyn, 1962). From crude polysaccharide, the only fraction eluted with 1 M NaCl (Fig. 2A) and UV –Visible spectrum of the EC polysaccharides exhibited typical characters of the polysaccharide. The UV-visible spectrum (Fig. 3A) of the polysaccharide did not show any absorption peaks at 260 nm and 280 nm which correlates with the absence of protein and nucleic acid respectively (Bian, Xie, & Chen, 2010). Previously, it has been reported that the presence of acidic polysaccharides with uronic acids and mucic acids in the leaves of *A. rosea, H.syriacus*, *C.olitorous* (Ohtani, Okai, Yamashita, Yuasa, & Misaki, 1995). Interestingly, similar compositions were observed with *E. caducifolia* as reported in roots of its family member *E. fisheriana* (Liu, Sun, Liu, & Yu, 2011). Acidic polysaccharides extracted from plant sources have been reported for their extensive biological activities like hypoglycemic, anticomplementary, immunomodulatory (Ohtani et al., 1995). Similar to that of other acidic polysaccharides, EC polysaccharides was also expected to show biological activities like antioxidant and immunomodulatory activities.

**Fig. 2.**
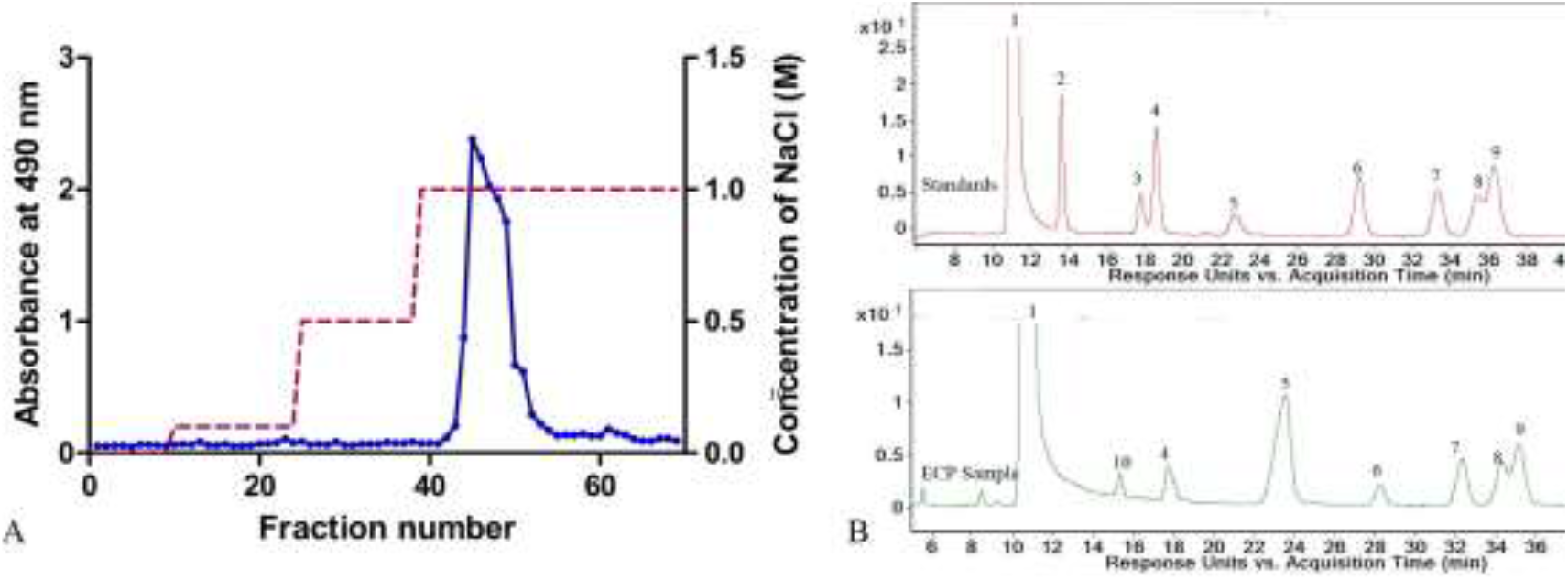
Chromatograms of the EC polysaccharide; A) Elution profile of EC polysaccharides purification on DEAE cellulose column, B) HPLC profile of the EC polysaccharide for monosaccharide analysis: 1.PMP, 2.Man, 3. GlcUA, 4.Rha, 5.Ino, 6. Glc, 7.Gal, 8.Xyl, 9.Ara, 10. GalUA.

**Fig. 3.**
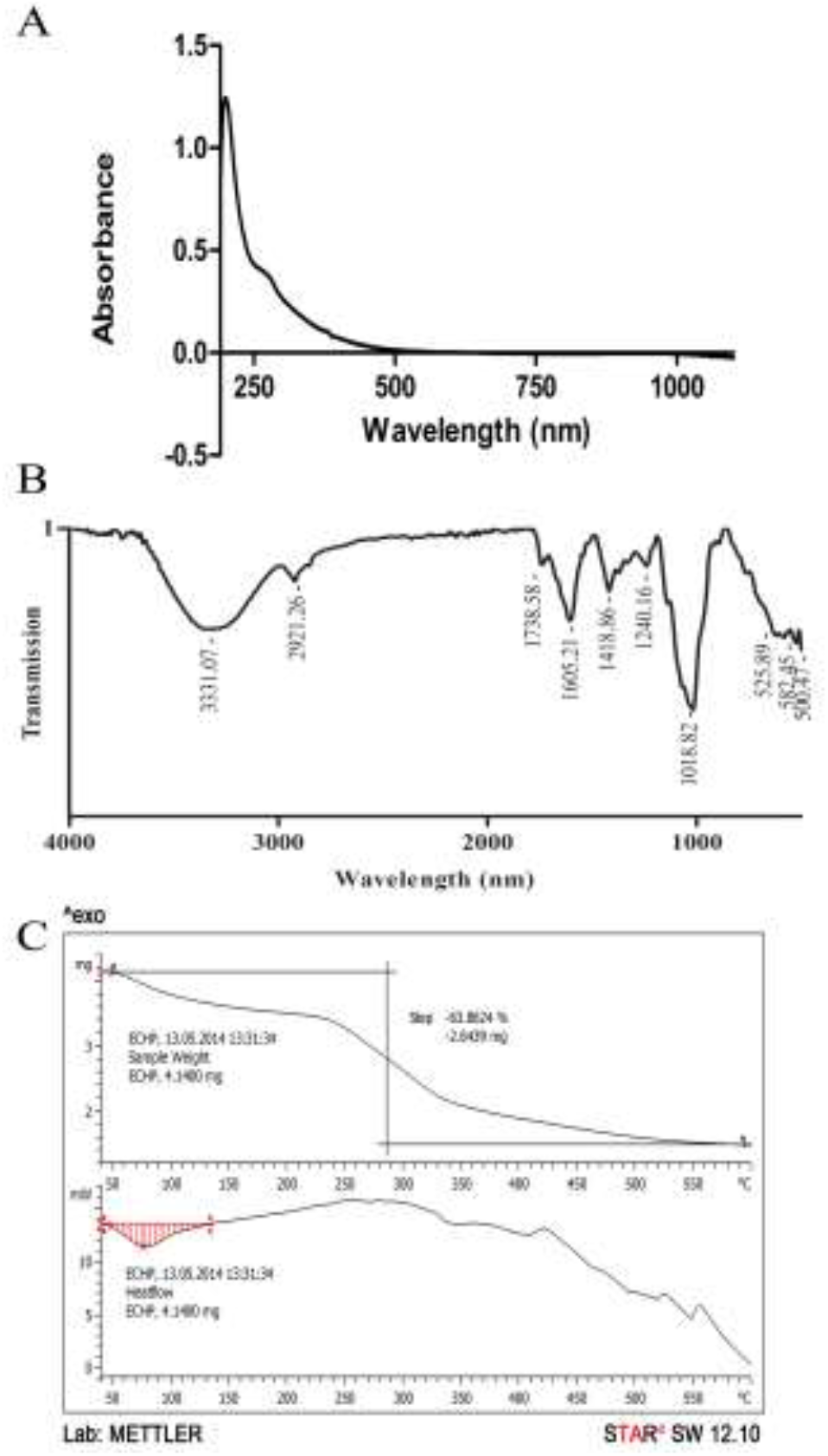
Spectra and thermogram of EC polysaccharides; A) UV-Visible Spectrum, B) ATR-IR Spectrum, C) TGDTA thermogram at heating rate 10^°^C min^-1^ under a nitrogen atmosphere.

### 3.3. Functional group analysis

Infrared spectra of the EC polysaccharides analysed to determine the functional groups present. Forbye the lack of sharp peaks at amine stretch in IR spectrum of the EC polysaccharides (Fig. 3B) mount the evidence that no contamination in purified polysaccharides. A characteristic broad peak at around 3330 cm^-1^ assigned to the hydroxyl group(-OH) present in the polysaccharide. An aliphatic C-H stretching vibrations at 2921 cm^-1^ and weak absorption at 1418 cm^-1^ is from aliphatic C-H bending of CH_2_ (Parikh & Madamwar, 2006). The two peaks towards the 1738 cm^-1^ and 1605 cm^-1^ in the spectra resulted from the COO^-^ deprotonated carboxylic group and two medium broad peaks at 1075 and 1018 cm^-1^ suggest the presence of C-O bonds. β-configuration of the monomers present in EC polysaccharides displayed with a peak around 800 cm^-1^ (Zhou, Zie, & Fu, 2000). Merged peaks in the fingerprint region indicate the linkages between monosaccharides.

### 3.4. Monosaccharide and Linkage analysis

Monosaccharide content of the polysaccharide was measured using PMP derivatization. Hydrolysed EC polysaccharides were derivatized with PMP and inositol was used as internal standard. Derivatised monomers were analysed with HPLC and confirmed the monosaccharide peaks with standards (Fig.2B). Molar contents were calculated according to Zhang et al., 2009 and noticed in the ratios of GlaUA, Rha, Glc, Gal, Xyl and Ara were in 1:2.5:4:3.6:3.7:3.3. *Euphorbia fischeriana*, a family member showed the high contents of Glucose and arabinose as compared with the *E. caducifolia* (Liu et al., 2011).

Methylated monosaccharides followed by silylation were determined by GCMS to analyse the linkage between the monomers present. From the chromatogram of the GCMS, it revealed the presence of 2,4 di (trimethyl silyl)-1,3,5,6 tetra--methyl galactouroran, 3,4 di (trimethyl silyl)-1,2,5,6 tetra-methyl galactouronan, 4,6 di (trimethyl silyl)-1,2,3,5, tetra-methyl galactouronan along with C2,C3,C4 positions of trimethyl silyl penta methyl galacturonans. Only two type of linkages were observed with rhamnose and arabinose:2,4 di trimethyl silyl-1,3,5,6 tetra methyl rhamnose and 2-tri methyl silyl-1,3,4,5,6 penta-methyl rhamnose and 2,5 di trimethylsilyl-1,3,4,6 tetra methyl arabinose, 2-trimethyl silyl-1,3,4,5,6 penta-methyl arabinose. Only linkage was noticed with glucose by forming 4-trimethylsilyl-1,2,3,5,6 methyl glucose. For galactose, 3,4 di trimethyl silyl-1,2,5,6 tetra methyl galactose and 4,6 di trimethyl silyl-1,2,3,5 tetra methyl galactose along with C2,C3,C4 positions of trimethyl silyl penta methyl galactoses. Xylose monomers showed 2,4 di trimethylsilyl-1,3,5,6 tetra methyl xylose, 2,3,4 tri tri methylsilyl 1,5,6 tri methyl xylose, and C2,C4 positions of trimethyl silyl penta methyl xylose were observed. Overall, it indicated that except glucose all monosaccharide are in the terminal positions, and possibility of arabinose, rhamnose involved in backbone formation along with xylose, galactoseand galacturonic acid.

### 3.5. Thermal analysis

Thermal nature of the polysaccharides was analyzed using TG-DTA. The thermogravimetric curve (Fig. 3C) of the EC Polysaccharides showed decomposition with three differentiated steps. The first mass loss event occurred from 50°C to 120°C with the loss of mass 10-15 % due to the diminution of adsorbed water which represent the hydrophilic nature of the polysaccharide (Zohuriaan & Shokrolahi, 2004). The Second stage starts at 230°C retains till 350°C and observed a loss of weight ca.36% at 285°C because of thermal depolymerization.The carbonization of polysaccharide causes sustained weight loss in the last stage occurred above 350°C to 600°C. The presence of charged cations like Na^+^, K^+^, Ca^2+^ and complex configuration of the polysaccharide resist the degradation process and left the solid residue around 35% after 600°C (Cerqueira et al., 2011; De, Ruiz-bermejo, Menor-salván, & Osuna-esteban, 2011). DTA thermogram of the EC polysaccharide showed early endotherm peak at 78°C with an onset temperature 48°C and end set point 114°C. Enthalpy change (ΔH) of 135.15 J/g was noticed at early stages. No glass transition (T_g_) temperature was exhibited in the thermogram which attributed to interference of early endothermic peak (Parikh & Madamwar, 2006).

### 3.6. Antioxidant Activity

A simple and sensitive, DPPH free radical scavenging activity was performed to observe the antioxidant activity of polysaccharides. Antioxidant activity was increased as the concentration of the polysaccharides increases and Ascorbic acid used as a standard to compare EC polysaccharides. Free radical scavenging activity of the polysaccharide gained momentum of activity after 0.5 mg/ml concentration to greater 35% to 65% (Fig. 4A). Besides DPPH, Nitric Oxide (NO^·^) free radical scavenging activity was also measured (Fig. 4B). NO^·^ react with oxygen to give free radicals nitrite and proxy nitrite, the reason for many inflammatory processes and cause of several diseases (Luyen et al., 2014). EC Polysaccharides, a competitor for oxygen to react with NO^·^ and thus inhibits the production of nitrates and nitrites. EC polysaccharides at 2 mg/ml concentration showed the free radical scavenging activity 65% with DPPH and 55% with Nitric Oxide. The ability of the polysaccharides to scavenge the free radicals in the reaction mixture more than 50% suggests as prospective antioxidants.

**Fig. 4.**
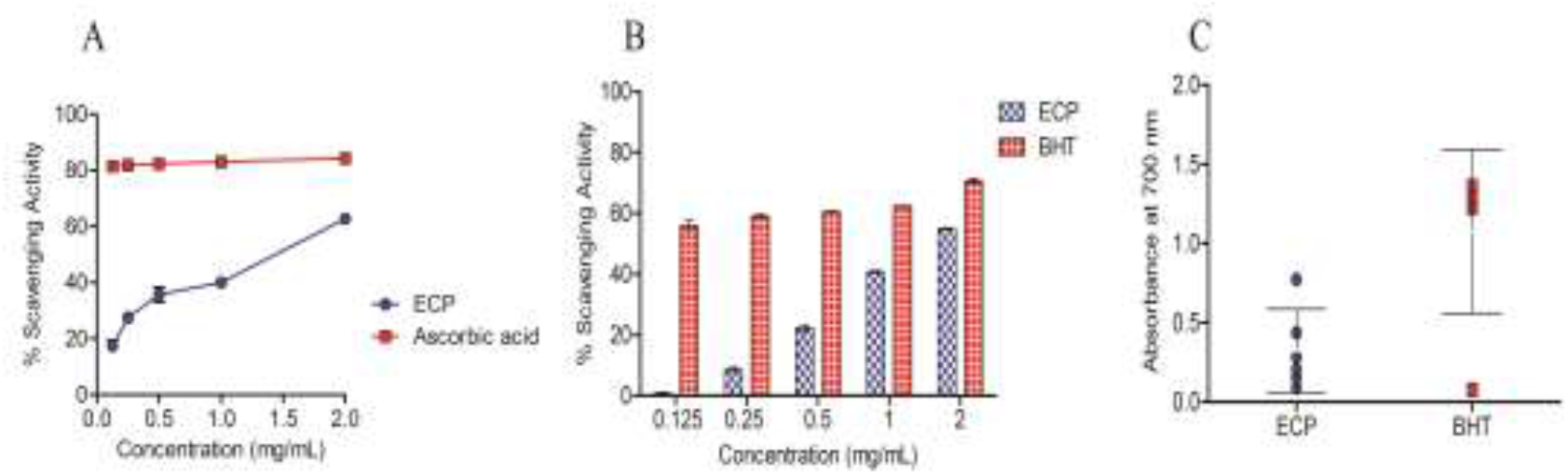
Antioxidant activity of EC polysaccharides; A) DPPH free radical scavenging activity, B) Nitric oxide radical scavenging activity, and C) Reducing power. (values are mean ± SE, n=3)

The reducing ability of a compound to donate electron can serve as a potential antioxidant. Apart from free radical scavenging activity, the antioxidant activity of a compound can also be evaluated by its capacity to reduce the ferric cyanide complex (Fe^3+^) to ferrous cyanide form (Fe^2+^). Thereby the colour change in the reaction mixture to different shades of green to blue indicates the antioxidant activity (Irshad, Zafaryab, Singh, & Rizvi, 2012). Similarly, green to blue shades were observed with EC polysaccharides, and prussian blue with BHT, where prussian blue indicates strong reducing activity. The absorbance was measured, as it proportionately correlates with the reducing power. Even though polysaccharide showed less degree of absorbance compared to BHT (Fig.4C), its reducing ability as an electron donor evinced with increasing polysaccharide concentration.

### 3.7. Emulsifying Activity

Emulsions are the mixture of immiscible liquids where droplets of one liquid dispersed in other with the help of an emulsifier. Most of the acidic polysaccharides like gum arabic, gum karaya, gum tragacanth are using as emulsifiers in the pharmaceutical and food industries (Benhura & Chidewe, 2004). In an interest to that, the efficacy of the EC polysaccharide as an emulsifying agent was tested by turbidimetry method. The emulsification ability of the EC polysaccharide was observed under a microscope, further measured the turbidity with a spectrophotometer (Fig. 5). The undiluted emulsion showed flocculation of droplets into various sizes. Further dilution with 0.1% sodium dodecyl sulfate, resulted in more dispersed globules and aggregates of floccules on standing for 24 h. In turbidimetry, increasing the concentration of the polysaccharide to oil shown linearity of the emulsification, and observed 36% of turbidity shift at high concentration and 88% of shift at lower concentrations of polysaccharide on long standing. Stabilization of dispersed phase in continuous phase even after 24 h suggests that the EC polysaccharide as a good emulsifier. Phase separation of the liquids occurred within 1 hour in the control where no polysaccharide was added.

**Fig. 5.**
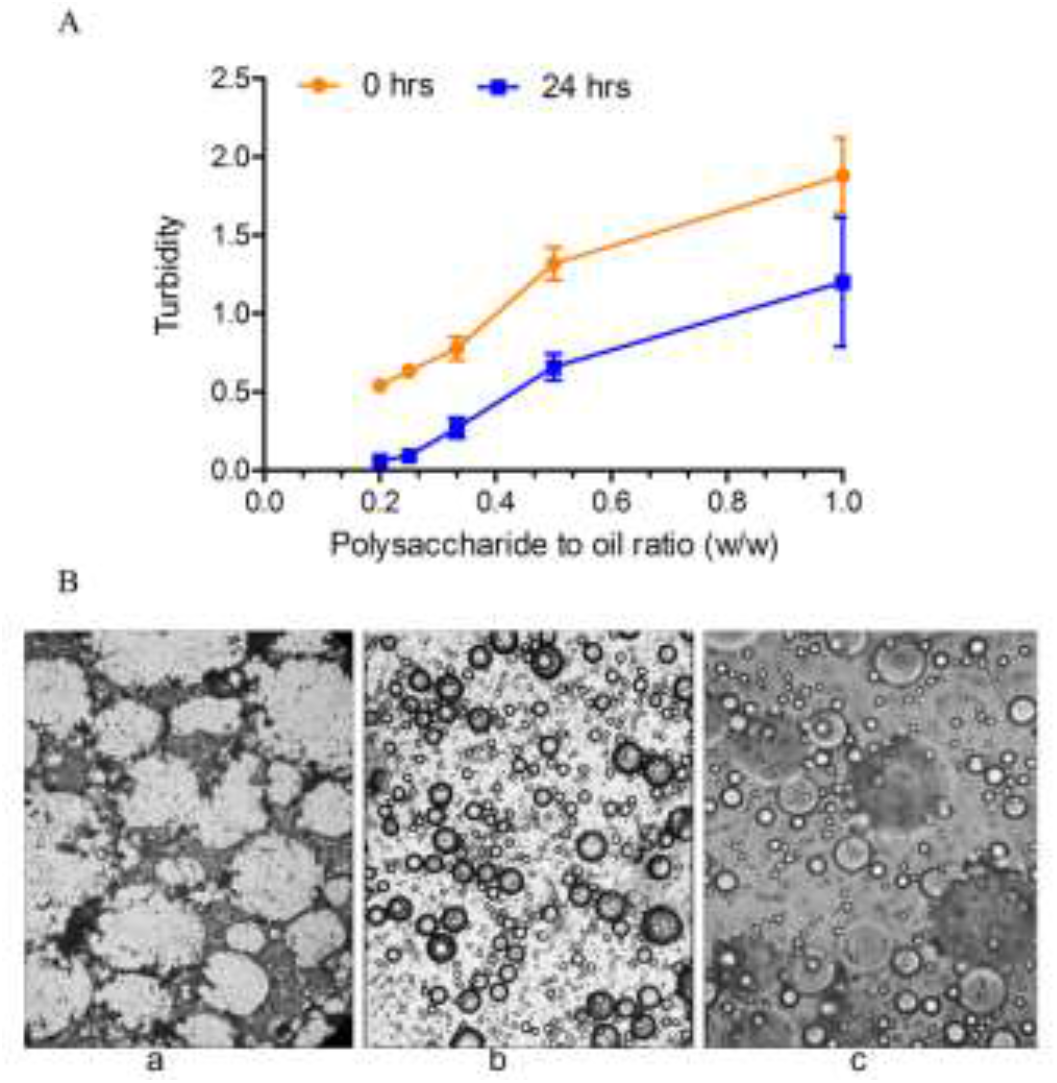
Emulsion ability of the EC polysaccharides; A) The turbidity of emulsion measured at 500 nm at 0 h and 24 h, B) Photomicrograph of Emulsion: a) Control (without polysaccharide), b) Emulsion at 0 h, c)) Emulsion at 24 h. (Values are mean±SE, n=3)

### 3.8. MTT Assay and Cytokine measurement

Polysaccharides from most of the plant sources are non-toxic in nature and modulate cytokine and chemokine production as an immunomodulator (Schepetkin & Quinn, 2006). Non-toxicity and immunomodulatory activities place the polysaccharides as ideal therapeutic candidates. The EC polysaccharides showed non-toxic nature and also directly stimulate the proliferation of RAW 264.7 cells (Fig. 6A). The proliferation effect was observed in the range of 50 ng/mL-1 μg/mL, i.e., from 107% to 150%, a further increase in the concentration depicted no significant cytotoxicity on cells compared to untreated cells.

**Fig. 6.**
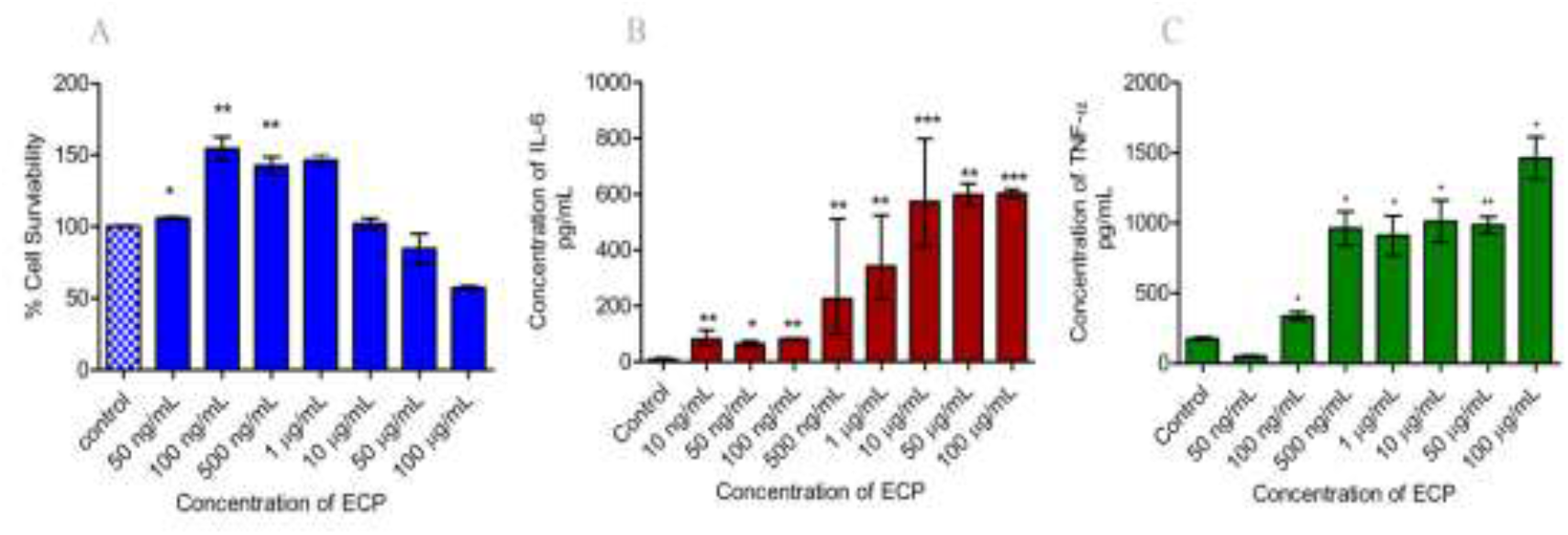
Effect of the EC polysaccharides on the RAW 264.7 at different concentration; A) Cell viability, and B) Concentration of the IL-6, C) TNF-α measured by ELISA in the culture supernatant. (n=3, values= mean ± SE, * =P<0.05, **=P<0.01, ***=P<0.001)

Most of the botanical polysaccharides activate several signal transduction pathways of macrophages and stimulate Nitric oxide (NO), Reactive oxygen species (ROS), and production of inflammatory (IL-1, IL-6, IL-12) as well as anti-inflammatory cytokines (IL-10) along with chemokines (TNF-α) (Schepetkin & Quinn, 2006). Treatment of macrophages (RAW 264.7 cells) with various concentrations of EC polysaccharides significantly enhanced the production of the pro-inflammatory cytokines such as IL-6 and TNF-α whereas untreated macrophages produced negligible amounts (Fig. 6B, 6C). The maximum IL-6 production observed with dose effective from 500 ng/mL to 100 μg/mL. Cell proliferation observed with EC polysaccharides at lower concentrations may be due to the initial production of IL-6, which stimulates the extracellular signal-regulated kinases and Mitogen-activated protein kinase pathways (Ogata et al., 1997). Increased levels of TNF-α and IL-6 at higher concentrations might be responsible for the certain level of cell death (<40%). These results evident that the EC polysaccharides at lower concentrations can be used as a therapeutic agent for promoting wound healing and regeneration activities.

## 4. Conclusions

Altogether this study suggests that manipulation of the temperature influence the polysaccharide yield along with solid to liquid ratio and time. Polysaccharides extracted to a maximum yield of 2.3% at 120°C, and the optimum conditions were identified using the response surface methodology. Purified acidic polysaccharides with uronic acids shown typical nature of spectral as well as thermal characteristics. EC Polysaccharides act as potential antioxidants and as a good emulsifier. Moreover, polysaccharides exhibited immunomodulatory activity with non-toxic nature on RAW 264.7 cells.

## Acknowledgements

Kusuma Venumadhav acknowledges to Rajiv Gandhi National Fellowship, University Grant Commission (UGC), New Delhi, India for providing financial assistance as a Senior Research Fellow. Authors gratefully acknowledges B.Devender, Metabolomics facility, UH for analytica support in GC-MS and LC-MS.

## References

A Pettolino, F., Cherie, W., B Fincher, G., & Antony, B. (2012). Determining the polysaccharide composition of plant cell walls. Nature Protocols, 7, 1590–1607.

Afza, N., Khan, A. Q., Malik, A., & Badar, Y. (1989). Cyclocaducinol, a cycloartane type triterpene from Euphorbia caducifolia. Phytochemistry, 28(7), 1982–1984.

Benhura, M. A. N., & Chidewe, C. K. (2004). The emulsifying properties of a polysaccharide isolated from the fruit of Cordia abyssinica. International Journal of Food Science and Technology, 39(5), 579–583. https://doi.org/10.1111/j.1365-2621.2004.00809.x

Bian, C., Xie, N., & Chen, F. (2010). Preparation of bioactive water-soluble pachyman hydrolyzed from sclerotial polysaccharides of Poria cocos by hydrolase. Polymer Journal, 42(3), 256–260. https://doi.org/10.1038/pj.2009.329

Cerqueira, M. A., Souza, B. W. S., Simões, J., Teixeira, J. A., Domingues, M. R. M., Coimbra, M. A., & Vicente, A. A. (2011). Structural and thermal characterization of galactomannans from non-conventional sources. Carbohydrate Polymers, 83(1), 179–185. https://doi.org/10.1016/j.carbpol.2010.07.036

Chaiklahan, R., Chirasuwan, N., Triratana, P., Loha, V., Tia, S., & Bunnag, B. (2013). Polysaccharide extraction from Spirulina sp. and its antioxidant capacity. International Journal of Biological Macromolecules, 58, 73–78. https://doi.org/10.1016/j.ijbiomac.2013.03.046

Chavre, B.. (2013). Botanicals used on the treatment of snakebite in some parts of Maharashtra. Indian Journal of Plant Sciences, 2(1), 52–54.

De, J. L., Ruiz-bermejo, M., Menor-salván, C., & Osuna-esteban, S. (2011). Thermal characterization of HCN polymers by TG-MS, TG, DTA and DSC methods. Polymer Degradation and Stability, 96(5), 943–948. https://doi.org/10.1016/j.polymdegradstab.2011.01.033

DuBois, M., Gilles, K. a., Hamilton, J. K., Rebers, P. a., & Smith, F. (1956). Colorimetric method for determination of sugars and related substances. Analytical Chemistry, 28(3), 350–356. https://doi.org/10.1021/ac60111a017

Goyal, M., Nagori, B. P., & Sasmal, D. (2012). Wound healing activity of latex of Euphorbia caducifolia. Journal of Ethnopharmacology, 144(3), 786–790. https://doi.org/10.1016/j.jep.2012.10.006

Green, L. C., Wagner, D. A., Glogowski, J., Skipper, P. L., Wishnok, J. S., & Tannenbaum, S. R. (1982). Analysis of nitrate, nitrite, and [15N]nitrate in biological fluids. Analytical Biochemistry, 126(1), 131–138. https://doi.org/10.1016/0003-2697(82)90118-X

Huang, S. Q., Li, J. W., Wang, Z., Pan, H. X., Che, J. X., & Ning, Z. X. (2010). Optimization of alkaline extraction of polysaccharides from ganoderma lucidum and their effect on immune function in mice. Molecules, 15(5), 3694–3708. https://doi.org/10.3390/molecules15053694

Irshad, M., Zafaryab, M., Singh, M., & Rizvi, M. M. a. (2012). Comparative Analysis of the Antioxidant Activity of Cassia fistula Extracts. International Journal of Medicinal Chemistry, 2012, 157125. https://doi.org/10.1155/2012/157125

Janarthanan, P., Zin Wan Yunus, W. M., & Ahmad, M. Bin. (2003). Thermal Behavior and Surface Morphology Studies on Polystyrene Grafted Sago Starch. Journal of Applied Polymer Science, 90(8), 2053–2058.https://doi.org/10.1002/app.12798

Jermyn, M. A. (1962). Chromatography of Acidic Polysaccharides on Deae-Cellulose. Australian Journal of Biological Sciences, 15(4), 787–792.

Kapoor, B., Mishra, R., Acharya, S., Lakhera, S., & Purohit, V. (2013). Antimicrobial screening of some herbal plants of the Rajasthan Desert: An Overview. Unique Journal of Engineering and Advanced Sciences, 1(1), 38–40.

Khan, N., Rahim, S. S., Boddupalli, C. S., Ghousunnissa, S., Padma, S., Pathak, N., … Mukhopadhyay, S. (2006). Hydrogen peroxide inhibits IL-12 p40 induction in macrophages by inhibiting c-rel translocation to the nucleus through activation of calmodulin protein. Blood, 107(4), 1513–1520. https://doi.org/10.1182/blood-2005-04-1707

Kumar, R. S., Sivakumar, T., Sunderam, R. S., Gupta, M., Mazumdar, U. K., Gomathi, P., … Kumar, K. A. (2005). Antioxidant and antimicrobial activities of Bauhinia racemosa L. stem bark. Brazilian Journal of Medical and Biological Research, 38(7), 1015–1024. https://doi.org/10.1590/S0100-879X2005000700004

Liu, J., Sun, Y., Liu, L., & Yu, C. (2011). The extraction process optimization and physicochemical properties of polysaccharides from the roots of Euphorbia fischeriana. International Journal of Biological Macromolecules, 49(3), 416–421. https://doi.org/10.1016/j.ijbiomac.2011.05.029

Luyen, B. T. T., Tai, B. H., Thao, N. P., Eun, K. J., Cha, J. Y., Xin, M. J., … Kim, Y. H. (2014). Anti-inflammatory components of Euphorbia humifusa Willd. Bioorganic and Medicinal Chemistry Letters, 24(8), 1895–1900. https://doi.org/10.1016/j.bmcl.2014.03.014

Molyneux, P. (2004). The use of the stable free radical diphenylpicryl-hydrazyl (DPPH) for estimating antioxidant activity. Songklanakarin J. Sci. Technol., 26(2), 211–219.

Ogata, a, Chauhan, D., Teoh, G., Treon, S. P., Urashima, M., Schlossman, R. L., & Anderson, K. C. (1997). IL-6 triggers cell growth via the Ras-dependent mitogen-activated protein kinase cascade. Journal of Immunology, 159, 2212–2221.

Ohtani, K., Okai, K., Yamashita, U., Yuasa, I., & Misaki, A. (1995). Characterization of an Acidic Polysaccharide Isolated from the Leaves of Corchorus olitorius (Moroheiya). Bioscience, Biotechnology, and Biochemistry, 59(3), 378–381. https://doi.org/10.1271/bbb.59.378

Parikh, A., & Madamwar, D. (2006). Partial characterization of extracellular polysaccharides from cyanobacteria. Bioresource Technology, 97(15), 1822–1827. https://doi.org/10.1016/j.biortech.2005.09.008

Ros, J. M., Laencina, J., Hellín, P., Jordán, M. J., Vila, R., & Rumpunen, K. (2004). Characterization of juice in fruits of different Chaenomeles species. LWT - Food Science and Technology, 37(3), 301–307. https://doi.org/10.1016/j.lwt.2003.09.005

Schepetkin, I. A., & Quinn, M. T. (2006). Botanical polysaccharides: Macrophage immunomodulation and therapeutic potential. International Immunopharmacology, 6(3), 317–333. https://doi.org/10.1016/j.intimp.2005.10.005

Schepetkin, I. A., Xie, G., Kirpotina, L. N., Klein, R. A., Jutila, M. A., & Quinn, M. T. (2008). Macrophage immunomodulatory activity of polysaccharides isolated from Opuntia polyacantha. International Immunopharmacology, 8(10), 1455–1466. https://doi.org/10.1016/j.intimp.2008.06.003

Shaik, P., Valli, S., Rajeswari, B., & Kumar, S. P. (2013). Potential of Euphorbia caducifolia Haines as a renewable source for biofuel. Indian Journal of Energy, 2(2), 99–107.

Wang, Z., Luo, D., & Ena, C. (2007). Optimization of polysaccharides extraction from Gynostemma pentaphyllum Makino using Uniform Design. Carbohydrate Polymers, 69(2), 311–317. https://doi.org/10.1016/j.carbpol.2006.10.013

Xu, Q., Yajima, T., Li, W., Saito, K., Ohshima, Y., & Yoshikai, Y. (2006). Levan (beta-2, 6-fructan), a major fraction of fermented soybean mucilage, displays immunostimulating properties via Toll-like receptor 4 signalling: induction of interleukin-12 production and suppression of T-helper type 2 response and immunoglobulin E p. Clinical and Experimental Allergy: Journal of the British Society for Allergy and Clinical Immunology, 36(1), 94–101. https://doi.org/10.1111/j.1365-2222.2006.02401.x

Yang, X., Zhao, Y., Wang, Q., Wang, H., & Mei, Q. (2005). Analysis of the monosaccharide components in Angelica polysaccharides by high performance liquid chromatography. Analytical Sciences: The International Journal of the Japan Society for Analytical Chemistry, 21(10), 1177–80. Retrieved from http://www.ncbi.nlm.nih.gov/pubmed/16270574

Ye, C.-L., & Lai, Y.-F. (2015). Optimization of Extraction Process and Antioxidant Activity of Polysaccharides from Leaves of Artemisia argyi Levl. et Vant. Journal of Food Processing and Preservation, 39(6), 1309–1317. https://doi.org/10.1111/jfpp.12349

Ye, D., Jiang, Z., Zheng, F., Wang, H., Zhang, Y., Gao, F., … Shi, G. (2015). Optimized extraction of polysaccharides from Grateloupia livida (Harv.) yamada and biological activities. Molecules, 20(9), 16817–16832. https://doi.org/10.3390/molecules200916817

Ye, H., Wang, K., Zhou, C., Liu, J., & Zeng, X. (2008). Purification, antitumor and antioxidant activities in vitro of polysaccharides from the brown seaweed Sargassum pallidum. Food Chemistry, 111, 428–432. https://doi.org/10.1016/j.foodchem.2008.04.012

Yildirim, A., Oktay, M., & Bilaloglu, V. (2001). The Antioxidant Activity of the Leaves of Cydonia vulgaris. Journal of Medicine (Cincinnati), 31, 23–27.

Yuan, X., Zeng, Y., Nie, K., Luo, D., & Wang, Z. (2015). Extraction Optimization, Characterization and Bioactivities of a Major Polysaccharide from Sargassum thunbergii. PLoS ONE, 10(12), 1–11. https://doi.org/10.1371/journal.pone.0144773

Zhou, P., Zie, M., & Fu, B. (2000). A review of the studies on the polysaccharide structure. Journal of Nanchang University (Natural Science), 25(2), 197–204.

Zohuriaan, M. J., & Shokrolahi, F. (2004). Thermal studies on natural and modified gums. Polymer Testing, 23(5), 575–579. https://doi.org/10.1016/j.polymertesting.2003.11.001

Zou, Y., Jiang, A., & Tian, M. (2015). Extraction optimization of antioxidant polysaccharides from Auricularia auricula fruiting bodies. Food Science and Technology, 35(3), 1–6. https://doi.org/10.1590/1678-457X.6712

